# Gene editing is suitable to treat GM1 Gangliosidosis: a proof-of-concept study

**DOI:** 10.1101/2022.04.17.488473

**Authors:** Delphine Leclerc, Louise Goujon, Sylvie Jaillard, Bénédicte Nouyou, Laurence Cluzeau, Léna Damaj, Christèle Dubourg, Amandine Etcheverry, Thierry Levade, Roseline Froissart, Stéphane Dréano, Xavier Guillory, Leif A Eriksson, Erika Launay, Frédéric Mouriaux, Marc-Antoine Belaud-Rotureau, Sylvie Odent, David Gilot

**Affiliations:** INSERM U1242, COSS, Univ Rennes, Rennes, 35042, France; CHU Rennes, Service de Génétique Clinique, Centre de Référence Maladies Rares CLAD-Ouest, FHU GenOMEDS, ERN ITHACA, Hôpital Sud, Rennes, France; INSERM, EHESP, IRSET - UMR_S, 1085, Université Rennes 1, Rennes, France; Service de Cytogénétique et Biologie Cellulaire, CHU Rennes, Rennes, France; Department of Pediatrics, Competence Center of Inherited Metabolic Disorders, Rennes Hospital, Rennes, France; Laboratoire de Génétique Moléculaire et Génomique, Centre Hospitalier Universitaire de Rennes, Rennes; Univ Rennes, CNRS, IGDR (Institut de Génétique et Développement de Rennes), UMR 6290, ERL U1305, Rennes, France; Laboratoire de Biochimie, CHU de Toulouse, Pôle biologie, Institut Fédératif de Biologie, Toulouse; CHU Lyon HCL, LBMMS - Service Biochimie et Biologie Moléculaire, UM Pathologies Héréditaires du Métabolisme et du Globule Rouge, Bron; Univ Rennes, CNRS, ISCR (Institut des Sciences Chimiques de Rennes) - UMR 6226, Rennes, France; Department of Chemistry and Molecular Biology, University of Gothenburg, Göteborg, Sweden; Univ Rennes, CHU Rennes, Department of Ophthalmology, Rennes, France

**Keywords:** Base editing, patient, disease, gangliosidosis, lysosome

## Abstract

Ganglioside-monosialic acid (GM1) gangliosidosis, a rare autosomal recessive disorder, is frequently caused by deleterious single nucleotide variants (SNVs) in *GLB1* gene. These variants result in reduced β-galactosidase (β-gal) activity, leading to neurodegeneration associated with premature death. Currently, no effective therapy for GM1 gangliosidosis is available. Three ongoing clinical trials aim to deliver a functional copy of the *GLB1* gene to stop disease progression. Here, we show that 41% of *GLB1* pathogenic SNVs might be cured by adenine base editors (ABEs). Our results demonstrate that ABE efficiently corrects the pathogenic allele in patient-derived fibroblasts, restoring a therapeutic level of β-gal activity. Unbiased off-target DNA analysis did not detect off-target editing activity in treated patient’s cells except a bystander edit without consequences on β-gal activity. Altogether our results suggest that gene editing is an alternative strategy to cure GM1 gangliosidosis, by correcting the root cause of disease and avoiding repetitive adeno-associated virus injections.

## Introduction

During the two last decades, high throughput sequencing (HTS) technology has accelerated the identification of genetic alterations involved in human diseases, thus bringing an accurate diagnosis for each patient. Despite this historical breakthrough, a high number of diagnosed patients are currently awaiting personalized therapy. The development of curative gene therapies for these patients is indispensable^1^. According to ClinVar, ∼54,500 human genetic variants have been associated with diseases and more than 58% are single nucleotide variants (SNVs) (https://www.ncbi.nlm.nih.gov/clinvar/). This observation suggests that gene therapy strategies aimed at correcting the root cause of disease could be an alternative strategy to cure disease instead of bringing a novel functional copy of the altered genes ^2,3^.

Nowadays, gene editing approaches such as base and prime editing (BE and PE) enable specific restoration of a functional gene by correcting these deleterious variants ^3^. BE has been successfully used to cure different diseases in different models and species ^4–8^. To date, the most advanced application in humans is to cure sickle cell anemia via *ex vivo* targeting of haematopoietic stem and progenitor cells (HSPCs) from patients ^5^. BE also shows promise in limiting the deleterious effect of progerin (SNV altering the *Lamin A* mRNA splicing) as recently demonstrated in a murine progeria model ^8^ and to lower cholesterol in serum of primates by turning off the *PCSK9* gene ^7,9^.

Although this innovative gene therapy approach is promising, it is important to keep in mind the editing efficiency, the potential off-target effects and the challenge in reaching all targeted cells *in vivo* ^10,11^. To date, BE has the best efficiency rate compared to PE or CRISPR-Cas9 coupled to a donor. However, only C-to-T (CBE) and A-to-G (ABE) base editing methods are fully validated for gene therapy ^12,13^. Moreover, CBE and ABE are active on a wide editing window (max efficiency nucleotide 4 to 6 from the 5’ end of the sgRNA depending on the BE version), suggesting that bystander C or A nucleotides should be also modified into T or G within this editing window (bystander edits) ^14,15^.

To facilitate the design of sgRNAs for BE applications, several improvements have been implemented such as less stringent protospacer adjacent motif (PAM) than the canonical 5’-NGG-3’ (such as NRN and NYN PAM; SpRY Cas enzyme) and high fidelity (HF) BE Cas enzymes ^16,17^. So, the latest generation of BE are characterized by a high efficiency rate of editing and a low level of off-target edits since these Cas9 cleave only one DNA strand (nickase) and are sometimes HF, making them exploitable for gene therapy ^18^.

In humans, CBE and ABE should correct known pathogenic variants, 14 and 47% respectively ^3^. Here, we investigated whether BE may constitute an alternative strategy to cure GM1 gangliosidosis. Indeed, there are currently no effective therapies and only supportive treatments can be offered ^19^. GM1 gangliosidosis is an autosomal recessive, lysosomal storage disorder estimated to occur in 1 in 100,000 to 200,000 newborns ^20^. There are four different types of GM1 gangliosidosis based on age when symptoms first appear and severity of disease progression. Defects in the *GLB1* gene (coding the β-galactosidase (β-gal)) cause impaired enzyme activity leading to the toxic accumulation of gangliosides and neurodegeneration that presents as cognitive impairment, paralysis and early death. Three ongoing clinical trials (NCT03952637, NCT04273269, NCT04713475) aim to deliver a functional copy of the *GLB1* gene to slow down or stop disease progression, but cannot reverse damage already caused by the disease. For two clinical trials, the *GLB1* cDNA is carried by Adeno-associated viruses (AAV) injected into the cisterna magna to reach the neuronal cells *via* the cerebrospinal fluid and to break down GM1 ganglioside. Preclinical studies clearly demonstrated the ability of AAV to transport genes into neuronal cells, suggesting that these vectors should also bring both BE and sgRNA ^21–23^.

Here, we show that 82% of genetic alterations of *GLB1* gene are SNVs and 41% of pathogenic variants could be targeted by ABE. As a proof-of-concept experiment, we designed and validated *in vitro* an ABE strategy for a young patient with GM1 gangliosidosis. Based on the ABE efficiency and the quantification of off-targets edits, our data suggest that gene editing is an alternative strategy to cure GM1 gangliosidosis.

## Methods

### Editorial Policies and Ethical Considerations

Written informed consent was obtained from the patient and his parents. All procedures were in accordance with the ethical standards of the Ethics Committee of Rennes University Hospital and the French law (CCTIRS Comité Consultatif sur le Traitement de l’Information en matière de Recherche dans le domaine de la Santé). Peripheral blood samples (for DNA extraction) and skin biopsies (for functional analyses and personalized gene therapy development) were collected from all the participants.

### Exome Sequencing and bioinformatics pipeline

Trio exome sequencing was performed at Rennes Hospital University (Molecular Genetics and Genomics Laboratory). DNA was extracted from blood samples using the Hamilton automate machine. Exome DNA library was prepared with the Agilent Focused Exome preparation kit. High-throughput sequencing was performed on a NextSeq5500 sequencer (Illumina) with a 2×75 bp paired-end running method. The BWA-MEM algorithm ^24^ was used to map the reads on the reference genome (GRCh37/hg19). The variant calling was performed according to GATK ^25^ and FreeBayes best practices. The ANNOVAR ^26^ and ALAMUT (Interactive Biosoftware) tools were used for variant annotation. According to variant localization and nature, functional impact was predicted with dbNSFP v3.0 database ^27^, SpliceSiteFinder-like, MaxEntScan, NNSPLICE and GeneSplicer. Variant prioritization was performed according to these predictions and information available on OMIM and ClinVar databases. Identified pathogenic variants were described according to Human Genome Variation Society nomenclature guidelines (https://varnomen.hgvs.org/).

### Patient-derived fibroblasts: primary culture and immortalization

The fresh skin punch biopsies were dilacerated with scalpels under a sterile area and then transferred into T25 flasks containing 5 mL of AmnioMAX C-100 (Gibco) medium. After 1-2 weeks, explant growing cells were washed with Dulbecco’s phosphate-buffered saline (DPBS, Lonza), trypsinized with TrypLE™ Express Enzyme (1X) (Gibco) and passaged in T75 flasks for fibroblasts amplification. Primary cells were then immortalized by SV40 T antigen lentiviral transduction. Lentiviral particle production was performed in HEK293T cell line by following Trono Lab recommendations (http://tronolab.epfl.ch). Briefly, psPAX2 (Addgene ID plasmid #12260), pVSVG (Addgene ID #8454) and pLOX-Ttag-iresTK plasmids (Addgene ID Plasmid #12246) were transfected into HEK293T cells with lipofectamine 2000 (Thermo Fisher Scientific) according to the manufacturer’s instructions. Cell supernatant containing lentiviral particles was collected after 2 days, centrifuged and filtered (0.45µm) to eliminate HEK293T-derived cell debris. Fibroblasts in primary culture (P6 well plate, Falcon®) were infected with 500µL of this infectious media in the presence of 8µg/mL polybrene (Sigma) to achieve viral infection.

### Cell culture

Immortalized patient-derived fibroblasts and HEK293T (ATCC) cells were cultured respectively in RPMI-1640 medium (Lonza) or DMEM (Gibco), each supplemented with 10% fetal bovine serum (FBS) (Gibco), 1% glutamine (Lonza) and 1% penicillin-streptomycin (Sigma). When confluent, cells were washed with DPBS (Lonza), digested with TrypLE™ Express enzyme (1X) (Gibco) and subsequently passaged at 1:10 ratio. All cells were maintained at 37°C in a humidified incubator with 5% CO_2_. Cell lines were routinely tested for mycoplasma infection with PlasmoTest Mycoplasma Detection Kit (Invivogen) and negative results were obtained.

### DNA extraction, PCR and Sanger sequencing

DNA was extracted from frozen cells pellets with the Nucleospin tissue DNA extraction kit (Macherey-Nagel). DNA concentration was evaluated using the NanoDrop1000 spectrophotometer (Thermo Scientific). PCR was performed from 50ng DNA with the Phusion™ High-Fidelity DNA Polymerase kit (Thermo Scientific). The list of primers is present in Table S1. Samples were subjected to thermal cycling as followed: 98°C 30sec (initial denaturation step), 98°C 10sec, primers annealing temperature 15sec, 72°C 30sec (35 cycles) and 72°C 5min (final elongation step). Amplicon size and purity were checked on 1,5% agarose gel electrophoresis before Sanger’s reaction. Sanger sequencing was performed directly on 1µL of PCR reaction in the presence of 5µM primer and the Big Dye Terminator V3.1 (Applied Biosystems™). The following thermocycler program was used: 96°C 5min (initial denaturation step), then 96°C 1min, primers annealing temperature 1min and 62°C 2min (30 cycles). Sequence products were purified on Sephadex G50 beads (GE Healthcare) and directly loaded onto 3130xl Genetic Analyzer capillary electrophoresis laser coupled system (ABI PRISM).

### RNA extraction, reverse transcription and quantitative PCR

RNA extraction was performed from frozen tissue in culture plates (P6 well plate, Falcon®) by using the NucleoSpin RNA kit (Macherey-Nagel). RNA concentration was measured with NanoDrop1000 spectrophotometer (Thermo Scientific). Reverse transcription was performed on 500ng RNA with the High-Capacity cDNA Reverse Transcription kit (Applied Biosystems). Quantitative PCR (qPCR) was performed in 384 well plates on 2.5 ng of reverse-transcribed cDNA using the SYBR Green PCR Master Mix (Applied Biosystems) in the presence of 1µM of forward and 1µM of reverse primers, using the QuantStudio 5 quantitative PCR equipment (Applied Biosystems). The primer sequences used are available in Table S1. Raw data were extracted with QuantStudio Design and Analysis Software (Applied biosystems). Relative gene expression compared to control conditions was calculated by using the 2^-ΔΔ^ Ct method. *GAPDH* was used as a housekeeping gene for normalization.

### Protein extraction

Cells were lysed and proteins were extracted from fresh fibroblast cells with RIPA buffer (Thermo Scientific) supplemented with protease and phosphatase inhibitor mini-tablet (Pierce, Thermo Scientific). Protein quantification was performed with Pierce BCA protein assay kit (Thermo Scientific) and absorbance was measured with spark microplate reader (TECAN) at 562nm.

### β-galactosidase enzymatic activity measurement

β-galactosidase activity was evaluated using a fluorogenic method initially described by Ho and O’Brien ^28^. Briefly, 20µL of fresh cell lysates were incubated in a white 96 well microplate (Grenier bio-one) with 100µL of buffer solution (pH 4.3) containing 0.5 mM of 4-methylumbelliferyl-β-D-galactopyranoside substrate (Sigma). A kinetic analysis was performed at 37°C for 1-2 hours. Every 4 minutes, the fluorogenic substrate was excited at 366nm and fluorescence emission, proportional to enzyme activity, was measured at 442nm with a spark microplate reader (TECAN). Enzyme activity was assessed from the slope of the fluorescence = f (time) curve and normalized per µg of protein.

### Western Blot

Protein samples were reduced and denatured with lithium dodecyl sulfate and reducing agent 10 min at 70°C. Then, 20µg of each sample was loaded in 4 to 12% Bis-Tris Gel (NuPAGE, Invitrogen) and migration was done: 1h30, 200V, 400mA. Proteins were then transferred onto a nitrocellulose membrane (Invitrogen™ iBlot™ Transfer Stack) with iBlot transfer device system (invitrogen). After transfer and incubation with a blocking solution (1 hour, room temperature, TBS-Tween 0,1%; 5% Bovine Serum Albumin, Eurobio), the membrane was probed with the primary antibody overnight at 4°C (rabbit anti-beta galactosidase antibody, 15518-1-AP Protein Tech, dilution 1/1000: mouse anti-alpha-tubulin, T6199 Sigma-Aldrich, dilution 1/10 000) and then 1 hour at room temperature with the HRP linked secondary antibody (HRP-linked anti-rabbit IgG, #7074 Santa Cruz, dilution 1/2000; HRP-linked anti-mouse IgG, #7076 Santa Cruz, dilution 1/2000). Signal was detected with Amersham ECL select substrate (Cytiva) under a luminescent image analyzer (ImageQuant LAS 4000). Uncropped western-blots are available in FIG. S1.

### Cloning

Single guide RNAs were designed according to the recommendations of Walton RT *et al*. ^17^, Table S1. Designed oligonucleotides were integrated into BPK1520 backbone (Addgene plasmid #65777) by golden gate assembly as previously described^29^. Plasmids were transformed into NEB^®^ Stable competent *E. coli* C3040H (New England BioLabs) by following the manufacturers’ recommendations. Bacteria were seeded onto ampicillin (100µg/mL) LB agar plates and incubated overnight at 37°C. Isolated colonies were then amplified overnight at 37°C with LB growth medium and ampicillin (100µg/mL) under constant agitation. Plasmid DNA was purified with NucleoBond^®^ Xtra Maxi kit (Macherey Nagel) according to the manufacturers’ instructions. Constructs were verified by Sanger sequencing. The plasmid encoding the sgRNA n°8 used in this study is available on Addgene (BPK1520-sgRNA GLB1, Addgene #184378).

### ABE Transfection

Immortalized fibroblasts were seeded in 6 well plates (Falcon®) (500,000 cells/well). Approximately 24h after seeding (∼60% confluency), cells were transfected with 11.55µL of Viafect reagent (Promega) according to the manufacturer’s protocols with 2200ng of ABEmax(7.10)-SpRY-P2A-EGFP plasmid (Addgene Plasmid #140003) and 1100ng of sgRNA plasmid (addgene plasmid #65777). The medium was replaced 6 hours and 24 hours after transfection to eliminate dead cells. Forty-eight hours after transfection, cells were washed with DPBS (Lonza), digested with TrypLE™ Express enzyme (Gibco) and resuspended into RPMI-1640 medium (Lonza). GFP positives cells were sorted by FACS (FACSAria FUSION, Becton Dickinson). Sorted cells were either lysed for DNA extraction and high throughput sequencing or individualized by limit dilution for clonal amplification (96 well plates, Falcon®, 1 cell/well). Approximately 3 weeks after the limit dilution, pure clones were amplified in 12 well plates (Falcon^®^). To easily identify the edited cells, beta-galactosidase assays were performed and gene editing was confirmed by Sanger sequencing.

### siRNA transfection

Immortalized fibroblasts were seeded in 6 well plates (Falcon®; 500,000 cells/well). Approximately 24h after seeding (∼60% confluency), cells were transfected with 50nm siRNA CTRL or siRNA *GLB1* (IDT, sequence available in Table S1) with lipofectamine RNAimax (Thermo fisher scientific) according to the manufacturer’s instructions. Forty-eight hours after transfection, cells cultures were stored at −80°C and proteins were then extracted.

### Off-target identification (CRISPOR)

Off-target (OT) sites were predicted bioinformatically by using the CRISPOR online tool ^30^ (http://crispor.tefor.net). Putative OT sites were ranked according to cutting frequency determination (CFD) score ^31^ and top ten OT hits were selected for deep-sequencing (amplicons) (Data are available in Table S3). In addition, putative exonic OT sites were checked by exome sequencing.

### Amplicon high-throughput sequencing

Genomic regions of interest were amplified by PCR (Phusion™ High-Fidelity DNA Polymerase, Thermo Scientific or KAPA2G Robust HotStart PCR, Sigma) from genomic DNA samples. Amplicons were purified with AMPure XP magnetic beads (Beckman coulter). Then, the library was prepared with the SureSelect XT HS2 DNA kit (Agilent). We followed the manufacturer’s recommendations (except that initial DNA fragmentation, hybridization and capture steps were skipped). Pooled libraries were sequenced on a MiSeq instrument (Illumina) with a 2×250 bp paired-end running method (MiSeq Reagent Nano Kit v2 500 cycles, Illumina). The flow cell was loaded with 5 picomolar pooled libraries containing 5% PhiX control V3 (Illumina). Raw sequencing data were demultiplexed with Bcl2Fastq software (v2.19, Illumina) (data are available in Table S3).

### CRISPResso2

FastQ files were submitted to CRISPResso2 ^32^ for precise editing quantification. We used the following parameters: -wc=-10, -q=30, --min-bp-quality-or-N=30, --conversion-nuc-from=A, -- conversion-nuc-to=G. The matrix “selected-nucleotide-percentage-table-around-sgRNA” generated by CRISPRESSO was used to calculate the editing percentage. The line with nucleotide “N” (Q score < 30) was removed with a Python script (FIG. S2) to avoid editing underestimation (data are available in Table S4).

### 3D structure prediction

Maestro Software (Schrödinger Release 2021-3: Schrödinger, LLC, New York, NY, 2019) was used for visualization, basic molecular modelling steps (protein preparation, energy minimization, etc.), as well as for the molecular dynamic (MD) simulation trajectory using Desmond. Crystal structure of the GLB1 protein in complex with galactose (PDB ID: 3THC) served as the starting point for molecular modelling. The protein preparation wizard was used to prepare the crystal structure as follows: hydrogen atoms were added and possible metal binding states generated. The protonation and tautomeric states of Asp, Glu, Arg, Lys and His were adjusted to match a pH of 4.3 and possible orientations of Asn and Gln residues were generated. Hydrogen bond sampling with adjustment of active site water molecule orientation was performed using PROPKA. Water molecules with fewer than three hydrogen bonds to non-water molecules were deleted. Finally, the protein-ligand complexes were subjected to geometry refinements using the OPLS4 force field in restrained minimizations. The S302G mutant was then generated using the ‘Residue and Loop mutation’ tool included with maestro, followed by an energy minimization of the whole structure in implicit water. The GLB1 WT and S302G structures were then prepared for MD with the system builder panel using the following parameters: each system was solvated with TIP3P water under periodic boundary conditions with the minimum distance between any atom in the solute and the edge of the periodic box being 10.0 Å; Na+/Cl-counterions were added as appropriate for the neutralization of the system; 0.154 M NaCl. Molecular dynamic simulations for each system were performed with Desmond as follows: 3 × 100 ns total time; OPLS4 force-field; 100 ps trajectory recording intervals; system energy set to 1.2, NPT ensemble class; 300.0 K; 1.01325 bar; model relaxed before simulation. Quality control of our simulations was done using the ‘Simulation Quality Analysis’ tool as well as the ‘Simulation Interactions Diagram’ tool.

### Statistical analysis

All data are expressed as the mean – standard error of the mean of at least three individual experiments excepting the HTS experiments. Data were analyzed by Student’s t-test via GraphPad Prism v8.0.1 (GraphPad Software, San Diego, CA). p-Values were deemed to be statistically significant as follows: *p<0.05; **p<0.01;***p<0.001; ****p<0.0001. All raw data are available in Table S5.

## Results

The three ongoing clinical trials to cure GM1 gangliosidosis related to defects in the *GLB1* gene (which decrease or abolish the β-gal enzyme activity) are based on the addition of a codon optimized *GLB1* gene using AAV transduction of neuronal cells (NCT03952637, NCT04273269, NCT04713475). Transgenes transferred by this type of vectors are transcriptionally active and are maintained as an extra-chromosomal (episomal) form in the transduced cells for a long-term period ^22^. However, this strategy requires patients to receive several AAV injections during their life to maintain a therapeutic level of β-gal enzyme activity. So, we investigated here whether genome edition using ABE could be an alternative strategy to cure GM1 gangliosidosis by correcting *GLB1* point mutations, avoiding repetitive AAV injections into the cerebrospinal fluid (cisterna magna).

### *GLB1* genetic variants

Firstly, we analyzed the genetic alterations of the *GLB1* gene found in ClinVar to characterize the origin of GM1 gangliosidosis (FIG. 1 and Table S2). The *GLB1* variants are almost exclusively germline (>95%) single nucleotide variants (82%), the majority of which are missense (60%) and some frameshift (11%) (FIG. 1A-C). It is interesting to note that the *GLB1* noncoding sequence is also impacted (UTR and splice sites; ∼20%). Next, we focused on the 93 known pathogenic variants (class 5 corresponding to 17% of all variants, N=532) to estimate the ability of BE and PE to correct them (FIG. 1D, E). Importantly, PE, ABE and CBE should, in theory, correct 100, 41 and 15%, respectively, of *GBL1* pathogenic variants described in ClinVar (FIG. 1F). Since BE is currently the most efficient approach for gene editing in humans ^3^, we evaluated the ability of ABE to correct a missense variant associated with GM1 gangliosidosis as a proof-of-concept experiment. We selected a patient with a missense variant targetable by ABE (FIG. 2).

**FIG. 1:**
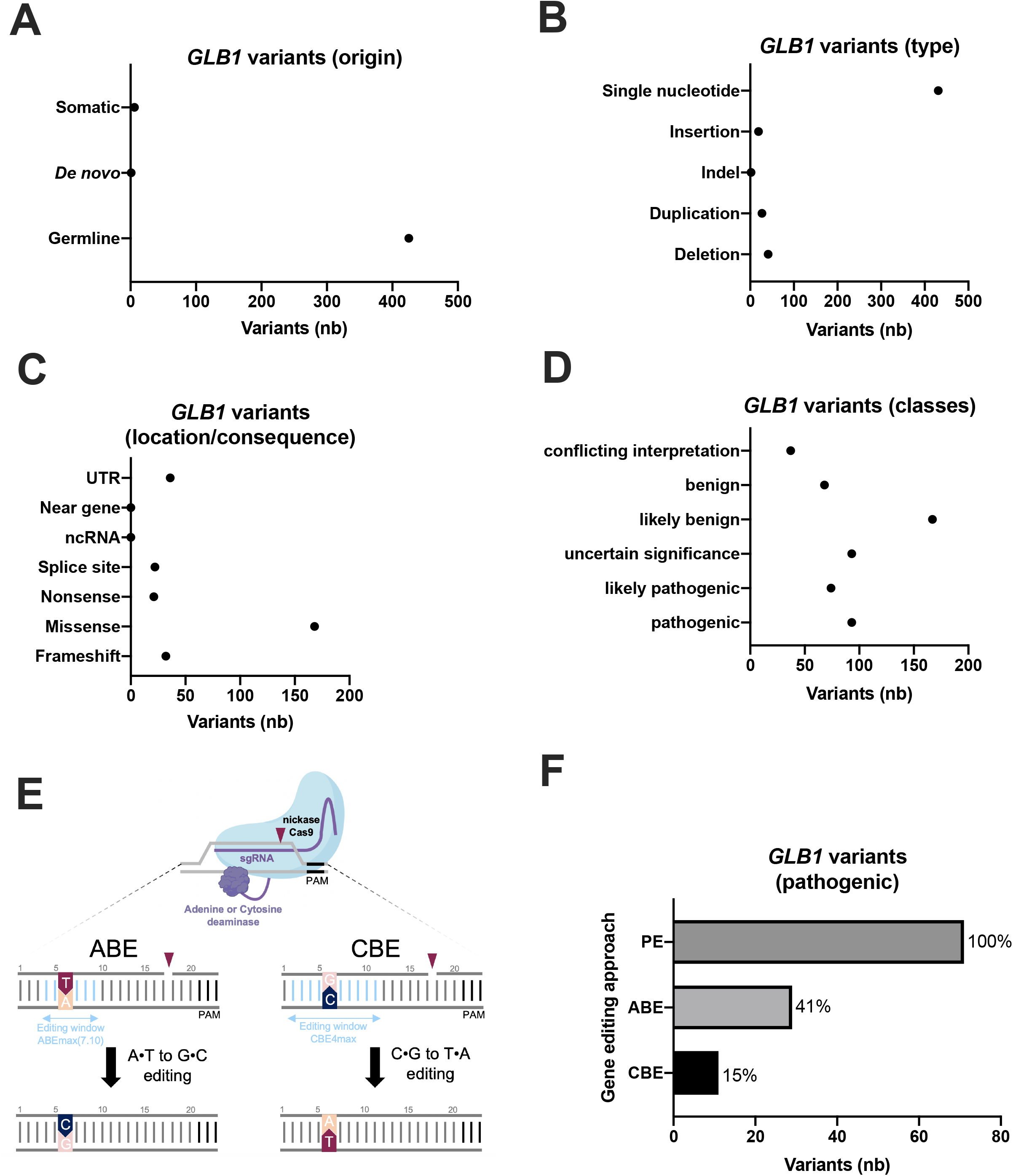
Genetic alterations of *GLB1*. **(A)** Origin of the *GLB1* variants (somatic, *de novo* or germline), (**B)** types of *GLB1* variants, (**C)** location and consequence of *GLB1* variants, **(D)** number of *GLB1* variants in the 5 classes, **(E)** scheme recapitulating base editor effects (ABE and CBE), **(F)** number of *GLB1* variants (class 5) that might be edited by PE (Prime Editing), ABE (Adenine base editor) or CBE (Cytosine base editor).

**FIG. 2:**
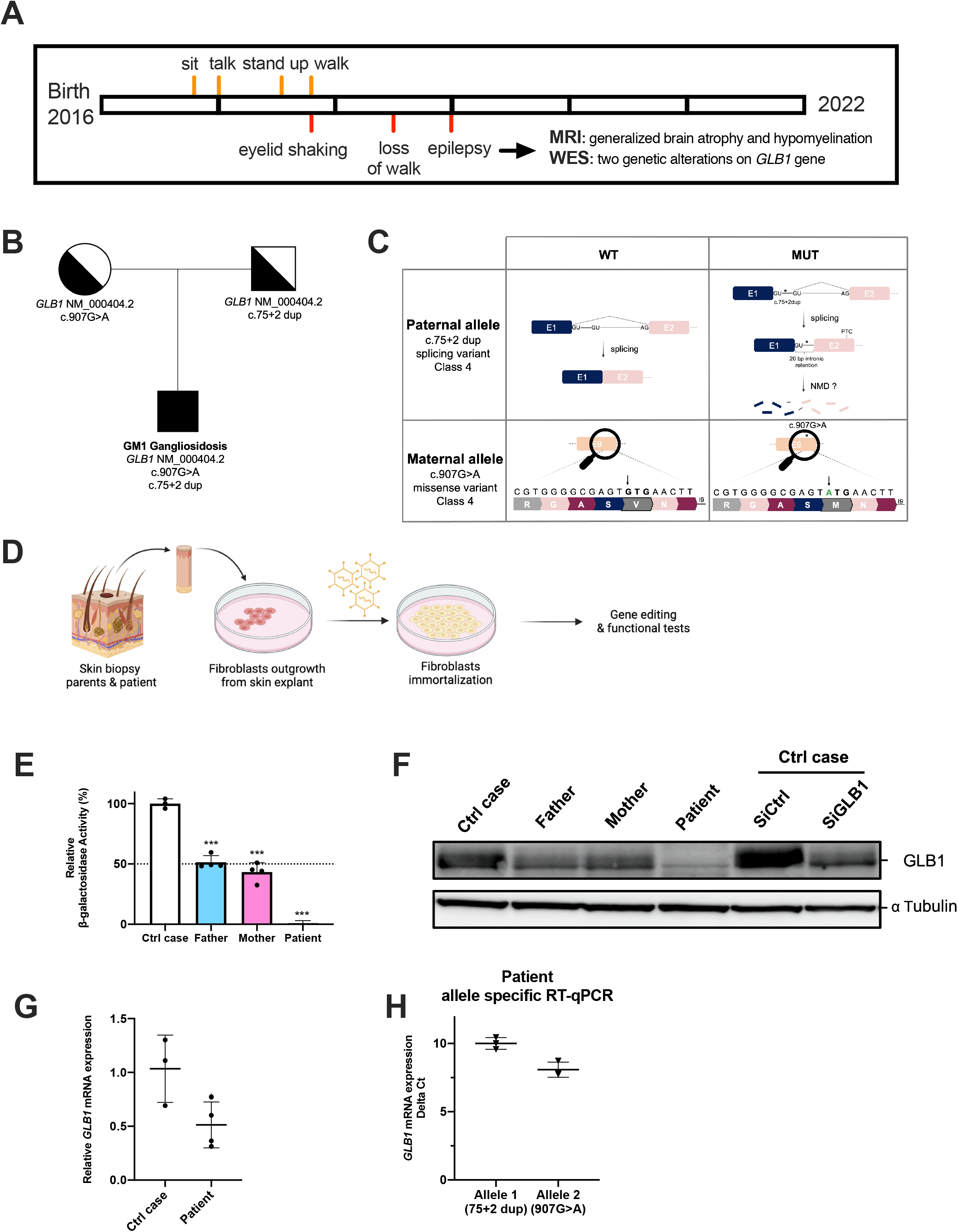
Characterization of patient-derived fibroblasts. **(A)** Milestones of the patient with GM1 gangliosidosis, (**B**) GM1 gangliosidosis patient pedigree, indicating autosomal recessive inheritance (genealogical tree), (**C**) hypothetical effect of inherited variants on *GLB1* mRNA and amino-acid sequence, (**D**) workflow describing patient-derived fibroblast purification and immortalization using SV40 T antigen (transduced by lentivirus), (**E**) β-galactosidase activity in patient-derived fibroblasts. Values were compared to values obtained from his parents and a control case (control fibroblasts), n≥3 biologically independent experiments, each histogram represents the mean ±s.d., Data were subjected to Student test *p<0.05; **p<0.01;***p<0.001; ****p<0.0001. (F) GLB1 protein levels in the same fibroblasts. Specificity of the GLB1 antibody raised against GLB1 protein was evaluated using siRNA targeting *GLB1* mRNA or a non-targeting siRNA (siCTR). Western blot results are representative of at least three experiments and alpha-tubulin serves as a loading control, (G) expression levels of *GLB1* mRNA evaluated by RT-qPCR in patient-derived fibroblasts or control fibroblasts, (H) allele specific qPCR quantification in patient-derived fibroblasts (delta Ct). NMD for nonsense mediated mRNA decay. Raw data are available in Table S4.

### *GLB1* characteristics of the patient

The patient is the second child of non-consanguineous parents (FIG. 2A). He has one healthy brother. Pregnancy and delivery were uneventful. He was born at 40 weeks of gestation, his birth weight was 4015 g. He achieved motor skills with slight delay, as he was able to sit at 9 months, to stand up at 19 months, and to make his first steps at 22 months of age (FIG. 2A). At the age of 30 months, he began to lose previously acquired milestones as he could no longer walk independently. Regarding speech development, he could pronounce his first words at the end of his first year of age, but he did not progress further and progressively lost speech during the following months. At the age of 22 months the parents noticed some episodes of eyelid shaking. The ophthalmologic examination was normal. A first brain magnetic resonance imaging (MRI) was performed at 23 months of age showing normal results. At the age of 3 years, generalized epilepsy became obvious. Initial anticonvulsant therapy by Levetiracetam was switched to valproic acid due to side effects. Follow-up cerebral MRI at the age of 3 years showed generalized brain atrophy and hypomyelination.

The first genetic investigations, which included screening for Steinert disease, Angelman and Prader Willi syndromes and CGH array analysis, were normal (data not shown). Trio whole exome sequencing of the child and his parents identified two heterozygous probably pathogenic variants in the *GLB1* gene (NM_000404.2): c.907G>A (p.Val303Met) and c.75+2 dup, inherited from his mother and father, respectively (FIG. 2B and 2C). The first variant is a missense variant (never described) and the second is located at a splice site, leading to a 20 nucleotide intron retention in *GLB1* RNA ^33^.

Since the *GLB1* genetic alterations suggested an altered β-gal activity associated with GM1 gangliosidosis, we evaluated it in white blood cells. Analyses revealed a pathogenic drop of β-gal activity (8 nmol/hour/mg of proteins in the patient versus control: 197 nmol/hour/mg of proteins, data not shown). The residual β-gal activity was estimated to be 4% of control. As a control, the neuraminidase activity was similar for the patient and the control sample (data not shown). Thus, the abnormal β-gal activity supported the pathogenicity of the two variants identified in *GLB1* and led to the diagnosis of a late-onset form of infantile GM1 gangliosidosis for this patient.

To confirm this diagnosis, three skin biopsies were performed to generate primary fibroblast cultures (parents and patient) (FIG. 2D). To obtain enough cells, we immortalized these fibroblasts using T antigen from SV40 via a lentiviral transduction.

In accordance with results obtained with PBMCs, we confirmed that patient fibroblasts displayed an undetectable β-gal activity in our experimental conditions (FIG. 2E). In contrast, the β-gal activity detected in fibroblasts isolated from the parents was reduced by 50% when compared to control fibroblasts. These results demonstrated that the combination of these two inherited mutations abrogated the β-gal activity and expression (FIG. 2F). Next, we confirmed a reduced amount of *GLB1* mRNA in patient-derived fibroblasts when compared to control fibroblasts (FIG. 2G) and both alleles were detectable (FIG. 2H). Allele 1 (c:75+2dup) has reduced expression compared to allele 2, probably due to nonsense-mediated decay of transcript containing 20bp of intronic sequence as already suggested ^33^. In conclusion, we showed that the combination of these two genetic alterations explains this case of GM1-gangliosidosis.

### Adenine Base Editing strategy for GM1 gangliosidosis

To conceive a therapeutic solution for this patient, we selected the ABE approach to correct only one allele; the GLB1 c.907G>A (p.Val303Met) mutation (maternally inherited), hoping to restore a β-gal activity (50%) as found in his asymptomatic parents (FIG. 3A). The ABE strategy is known to reach a greater efficiency than PE, and ABE is already used in clinical trials ^3^. Only a PE approach has the potential to repair the other genetic alteration inherited from the father ^34^.

**FIG. 3:**
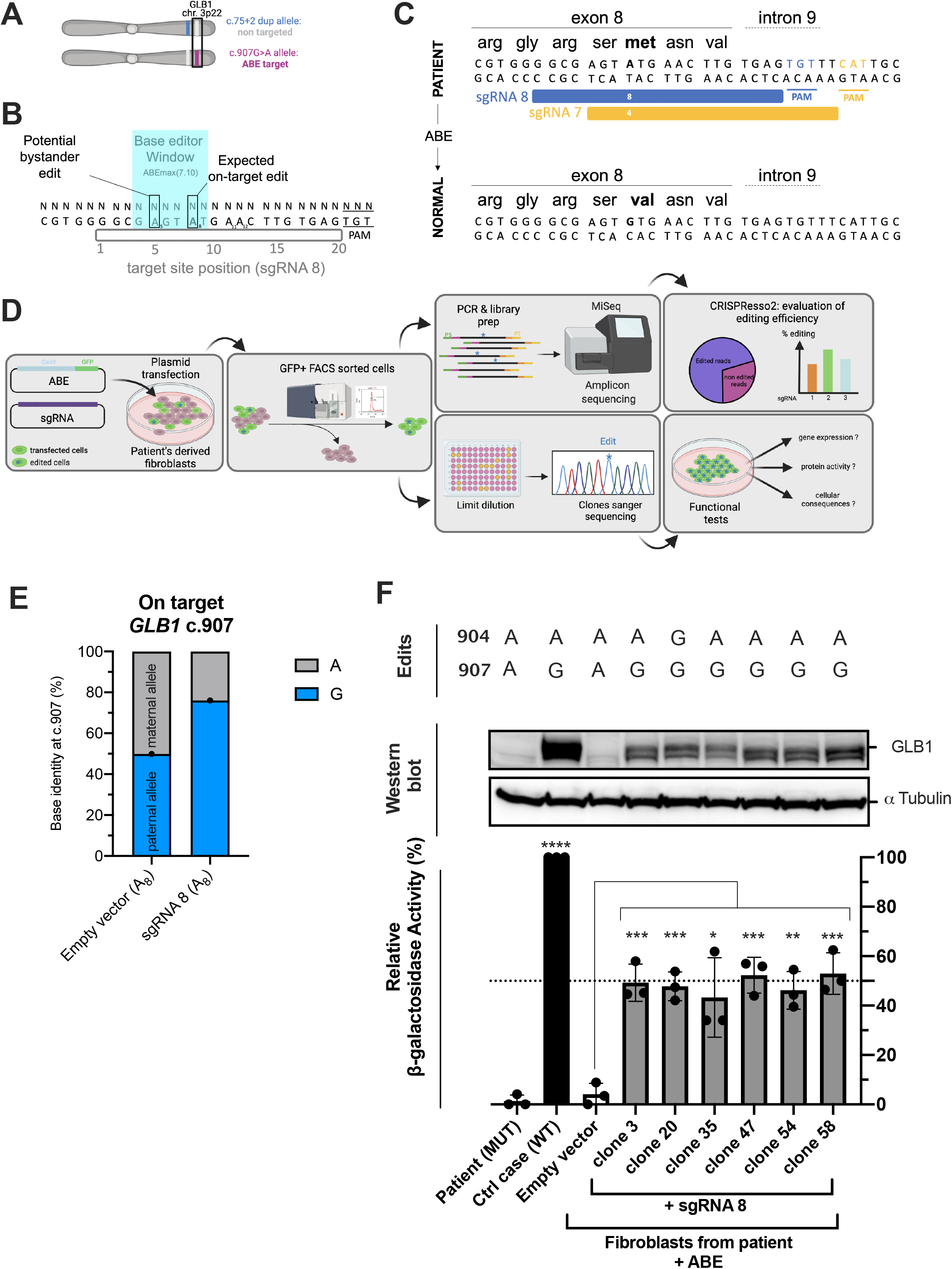
Adenine Base Editing strategy for GM1 gangliosidosis. **(A)** Maternal inherited *GLB1* variant is targetable by ABE, (**B**) scheme explaining the base editor window. Potential bystander edit might be obtained (proximal adenine to the on-target Adenine located in the base editor window), (**C**) two sgRNA were designed to correct the pathogenic ATG (Met) into the GTG (Val) on exon 8 of *GLB1* gene using adenine base editor, (**D**) workflow used to evaluate the ABE efficiency and specificity for the patient-derived fibroblasts, (**E**) efficiency of ABE to introduce the on-target edit (907A=>G) using the sgRNA8 in fibroblasts. Results were obtained by amplicon deep-sequencing and analyzed by CRISPResso2. Bystander edits were identified. (**F**) Clones isolated after transfection sgRNA8 + ABE or sgRNA CTR + ABE) were compared to normal fibroblasts or untreated patient-derived fibroblasts. Each clone was characterized for β-galactosidase activity, β-galactosidase protein and edits. WT for wild-type and MUT for mutated. n=3 biologically independent experiments, each histogram represents the mean ±s.d. Data were subjected to Student test *p<0.05; **p<0.01;***p<0.001; ****p<0.0001. Alpha-tubulin serves as a loading control. Raw data are available in Table S4

To optimize our chance of designing effective sgRNA for ABE, we selected the human codon optimized ABEmax(7.10): A-to-G base editor with nSpCas9 SpRY (D10A/L1111R/D1135V/G1218R/E1219F/A1322R/R1335V/T1337R) ^17^. Using this near-PAMless engineered CRISPR-Cas9 variant, we successfully designed and cloned two sgRNAs (FIG. 3A-C). Next, ABEmax(7.10) and one sgRNA were transfected into immortalized fibroblasts derived from the patient (FIG. 3D). Transfected cells were selected using the EGFP signal encoded by the plasmid pCMV-T7-ABEmax(7.10)-SpCas9-NG-P2A-EGFP (FIG. 3D). Five days after the enrichment of transfected cells (EGFP^+^ sorted cells), the BE efficiency was estimated by DNA-sequencing as previously described ^6,32^. CRISPResso2 analyses indicated that A907 was converted into G907 (reference nucleotide) in ∼52% of reads from the maternal allele as expected with this BE approach (FIG. 3E). The sgRNA 7 was inefficient (Table S4), confirming the requirement to evaluate several sgRNAs for BE.

Next, we performed a clonal selection to isolate edited clones: G907 (on-target edit). β-gal activity and β-gal protein levels were assessed (FIG. 3F). For the 6 clones exposed to the sgRNA8, ABE rescued a therapeutic level of β-gal protein and β-gal activity when compared to donor cells (wt *GLB1*). The restored β-gal activity reached a similar activity to that detected in the fibroblasts isolated from his asymptomatic parents (50% of the normal values) (FIG. 2E).

### Window editing of Adenine Base Editor

Base editors are wonderful tools to cure diseases but the bystander edits might limit their utility by modifying a proximal A (or C), and, in turn, giving an unwanted amino-acid (bystander edit). Closer inspection of our sequencing results indicated that A904 is also converted into G904 (∼27% of the reads) (FIG. 3F and 4A,B). This result could be explained by the active window of the ABE (FIG. 3B). Since the clone#20 (G907-G904; on-target and bystander edits) displayed a similar β-gal activity than other edited clones (G907-A904; only on-target edit), we investigated the neutral role of this unwanted edit.

**FIG. 4:**
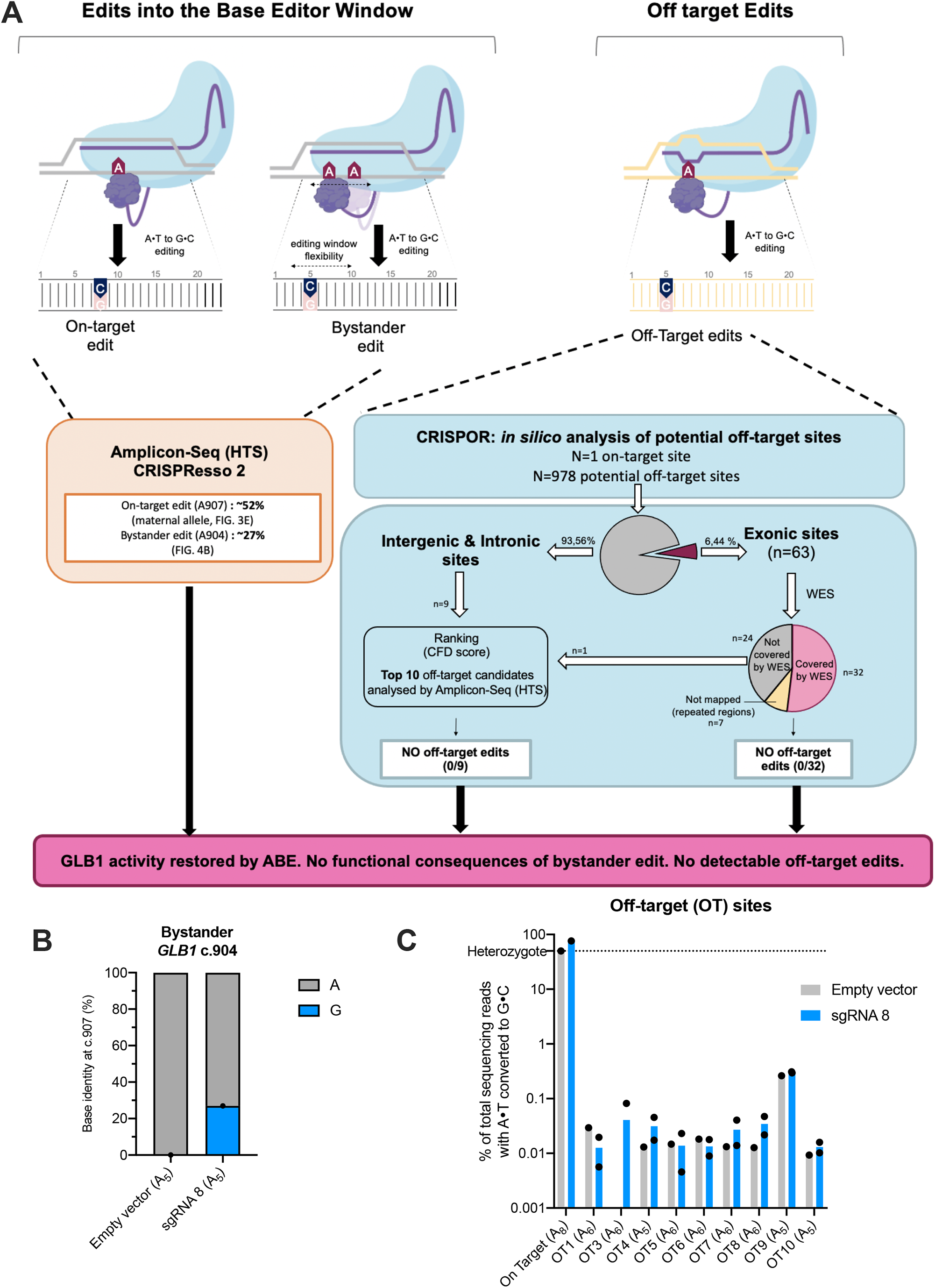
Specificity assessment of Adenine Base Editor for *GLB1* therapy. **(A)** Scheme recapitulating our workflow to identify the on-target and bystander edits on the *GLB1* gene (base editor window), (**B**) workflow explaining the off-target effects of ABE in patient-derived fibroblasts. Potential off-targets edits were predicted by CRISPOR and 41 potential sites were examined by combining whole exome sequencing and amplicon sequencing results. CFD for Cutting Frequency Determination score ^30,31^. (**C**) Analysis of the top10 off-target predicted sites for sgRNA8+ABE.

We predicted the 3D structure of GLB1 protein (FIG. S3). *In silico* mutation followed by an energy minimization of the whole structure showed minimal impact on the surrounding residues and GLB1 structure (RMSD_all-atoms_ = 0.720 Å). To study the potential impact of this mutation more in depth, molecular dynamics (MD) simulations were carried out (3 × 100 ns) on GLB1 wild type and S302G (FIG. S3A). Analysis of the trajectories did not reveal any significant changes in the secondary of tertiary structure of GLB1 (FIG. S3B). Altogether our results strongly suggest that this ABE off-target effect (bystander edit) will have no impact on GM1 gangliosidosis patients.

### Adenine Base Editor and off-targets detection

Base editors are known to generate bystander edits on surrounding A or C (window editing) (FIG. 3 & 4A). They can have negligible consequence as demonstrated here or avoided by targeting a region without modifiable surrounding bases. However, the off-target edits due to an imperfect sgRNA base-pairing to DNA might impair the clinical development of a sgRNA coupled to a base editor.

To uncover genomic loci with the potential for off-target editing, a computational prediction was used (CRISPOR ^30^) (FIG.4B). In total, 978 candidate sites have been predicted, including only 63 exonic sites (Table S3). Since it remains difficult to evaluate all these potential sites, we ranked them according to the Cutting Frequency Determination (CFD) score as previously published ^6^. We focused on the top-10 CFD scores (only one is located in an exon (*IDH1* gene, Table S3)) and off-target effects were investigated by deep-sequencing (amplicon-seq). Amplicons were analyzed *via* CRISPResso2 as previously done for BE efficiency measurement (FIG.3E). These samples were stringently quality filtered with a flag for minimum average read quality of 30 (phred33 score=30) to ensure SNP calling was only performed on high-quality reads. In our conditions, no off-target edits were found for the sgRNA8 in patient-derived fibroblasts. We failed to amplify the OT2 region by PCR. Thus, 9 among the top 10 CFD score sites were investigated (FIG. 4C). To reinforce this result, we performed whole exome sequencing (WES) (Clone #58 & #20 vs sgRNA control transfected cells). On the 63 exonic predicted off target sites, 32 sites were covered by WES. Under our conditions, no off-target edits were found for the sgRNA8 in patient-derived fibroblasts (0/32). In total, 41 potential off-target sites, including those with the highest CFD scores, were analyzed. We did not find any off-target effect using this protocol.

## Discussion

Gene therapy for GM1 gangliosidosis may be able to slow down or stop disease progression but cannot reverse damage already caused by the disease ^19,22^. There is currently no curative treatment for GM1 gangliosidosis. The three ongoing clinical trials aim at bringing a healthy *GLB1* coding sequence into the cells of the brain and spinal cord (AAV injections into the cisterna magna). To date, it is not possible to evaluate the number of nervous system cells that will be targeted by this classical gene therapy approach (efficiency). Patient follow-up is necessary to determine disease progression and potential adverse effects. Here, we provide a proof-of-concept study, strongly suggesting that genome editing can be an alternative strategy for this deleterious disease. To vehiculate the sgRNA and the ABE, the same AAV vector used for ongoing clinical trials ^22,23^ might be used for a single exposure. To obtain an optimal effect, the injection (genome editing) should be applied as soon as possible after birth, as already done for another neuronal disease using antisense oligonucleotides ^35^. However, viral delivery of base editors allows sustained expression in transduced cells, which may increase the frequency of off-target editing ^36,37^. In addition, viral vectors used in gene therapy raises the possibility of vector integration into the genome of patient cells, which may promote oncogenesis ^3,8,38^. Recently, another strategy has been published with Virus-like particles (VLPs) efficiently ‘infecting cells’ without carrying viral genetic material. So, the safety and efficiency of VLPs render them promising for gene editing approaches including *in vivo* ^7,39^.

### Limit of the study

Off-target edits might restrain the BE utility for human therapy. Here, by combining whole exome sequencing and amplicon sequencing for the 10 potential off-target sites ranked by CRISPOR, we did not identify off-target edits in our experimental conditions (0/41 sites). It is important to note that off-target editing independent of the DNA-sgRNA pairing has not been investigated here. For a clinical application, whole genome sequencing and RNA editing by base editor must be analyzed, according to the recent FDA press release (Human Gene Therapy Products Incorporating Human Genome Editing). Here, we performed a proof-of-concept study using patient-derived fibroblasts. Additional experiments are needed to replicate these results in cells derived from additional patients.

## Conclusion

GM1 gangliosidosis is a rare autosomal recessive disorder estimated to occur in 1 in 100,000 to 200,000 newborns. Here, we showed that 56% of the pathogenic mutations in *GLB1* gene might be targeted by the BE (ABE+CBE) and 100% by the PE. Interestingly, only one pathogenic allele needs to be corrected to cure the disease. Moreover, we demonstrated the efficiency and the safety of ABE in patient-derived fibroblasts in accordance with other BE studies ^8,16^. In conclusion, our study strongly suggests that gene editing is an alternative strategy to cure GM1 gangliosidosis immediately after the birth to limit irreversible damage.

## Acknowledgments

Firstly, authors would like to thanks the family (parents and son) for their kind support. Authors would like to thanks their collaborators for helpful advices, protocols and vectors: Pr C. Bendavid, Dr C. Moreau, Pr D. R. Liu (vectors), Pr B. Kleinstiver (pCMV-T7-ABEmax(7.10)-SpRY-P2A-EGFP (RTW5025)), the Biosit platforms especially Dr P. Gripon for BSL3 and L. Deleurme and A. Aimé for FACS analysis, the CRISP’edit platform (Bordeaux) especially Dr B. Turcq and Dr V. Prouzet-Mauleon, the head of the Inserm research unit Dr E. Chevet, the technicians from the Cell Biology service CHU Rennes and Pr E. Tucker for critical reading of this manuscript.

## Authorship contribution

Delphine Leclerc : Conceptualization, Data curation, Formal analysis, Investigation, Visualization, Validation, Writing – original draft, Writing – review & editing

Louise Goujon : Data curation, Investigation, Writing – review & editing

Sylvie Jaillard : Data curation, Formal analysis, Investigation, Project administration, Writing – review & editing

Bénédicte Nouyou: Data curation, Formal analysis, Validation, Software, Writing – review & editing

Laurence Cluzeau : Investigation, Writing – review & editing

Léna Damaj : Resources, Writing – review & editing

Christèle Dubourg : Data curation, Investigation, Resources, Writing – review & editing

Amandine Etcheverry : Data curation, Investigation, Writing – review & editing

Thierry Levade : Investigation, Resources, Writing – review & editing

Roseline Froissart : Investigation, Resources, Writing – review & editing

Stéphane Dréano : Data curation, Investigation, Writing – review & editing

Xavier Guillory : Investigation, Writing – review & editing

Leif A Eriksson : Investigation, Writing – review & editing

Erika Launay : Data curation, Investigation, Writing – review & editing

Frédéric Mouriaux : Project administration, Writing – review & editing

Marc-Antoine Belaud-Rotureau : Project administration, Writing – review & editing

Sylvie Odent : Resources, Project administration

David Gilot : Conceptualization, Data curation, Formal analysis, Investigation, Visualization, Validation, Writing – original draft, Writing – review & editing, Project administration.

## Supplementary material

Supplementary Table 1: sgRNA, pegRNA, primers sequences.

Supplementary Table 2: *GLB1* variants in ClinVar.

Supplementary Table 3: CRISPOR predictions for sgRNA 8 targeting *GLB1* gene.

Supplementary Table 4: CRISPResso2 results for On-target and off-targets.

Supplementary Table 5: Raw data

**Supplementary Figure 1:**
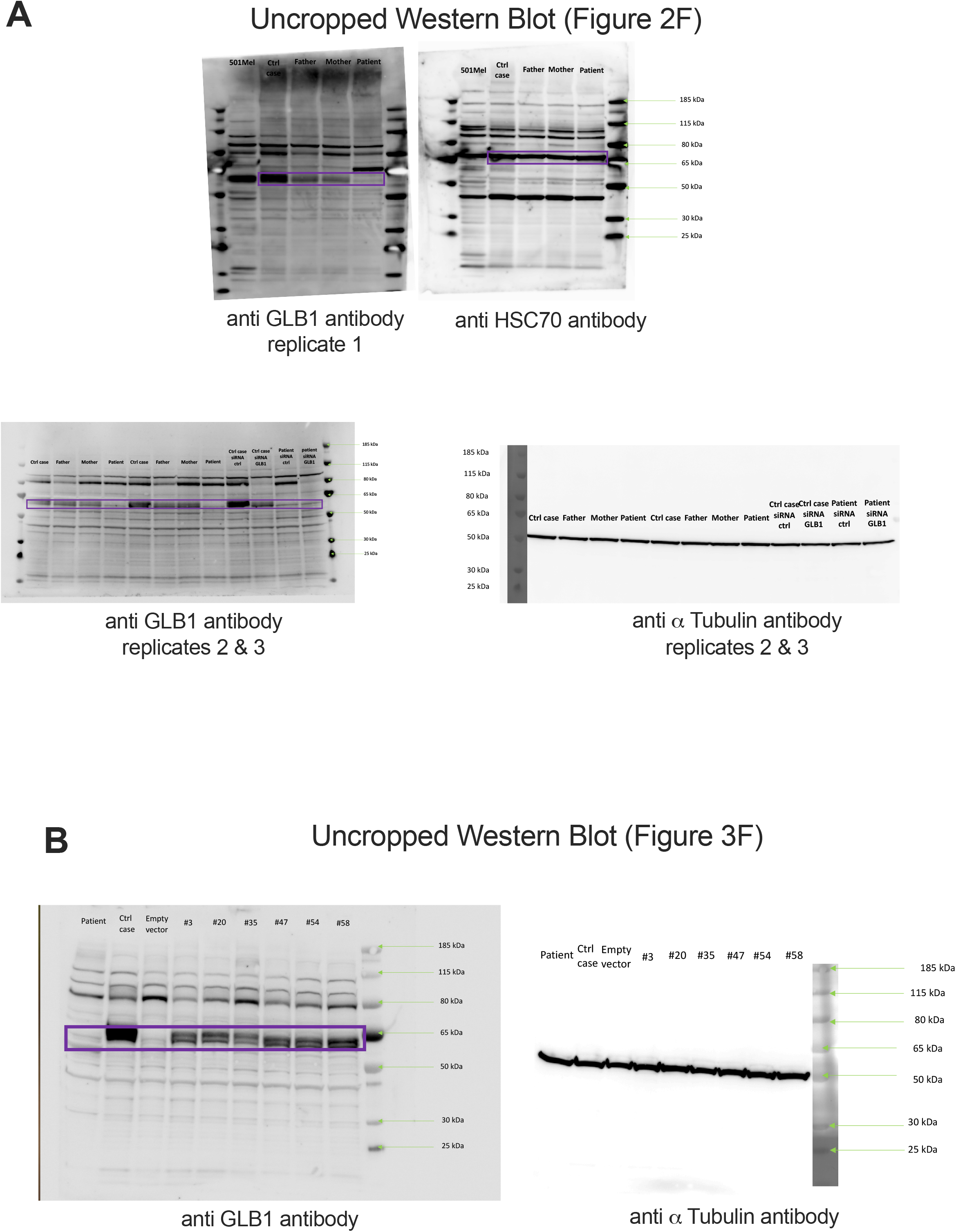
uncropped pictures of western-blot experiments.

**Supplementary Figure 2:**
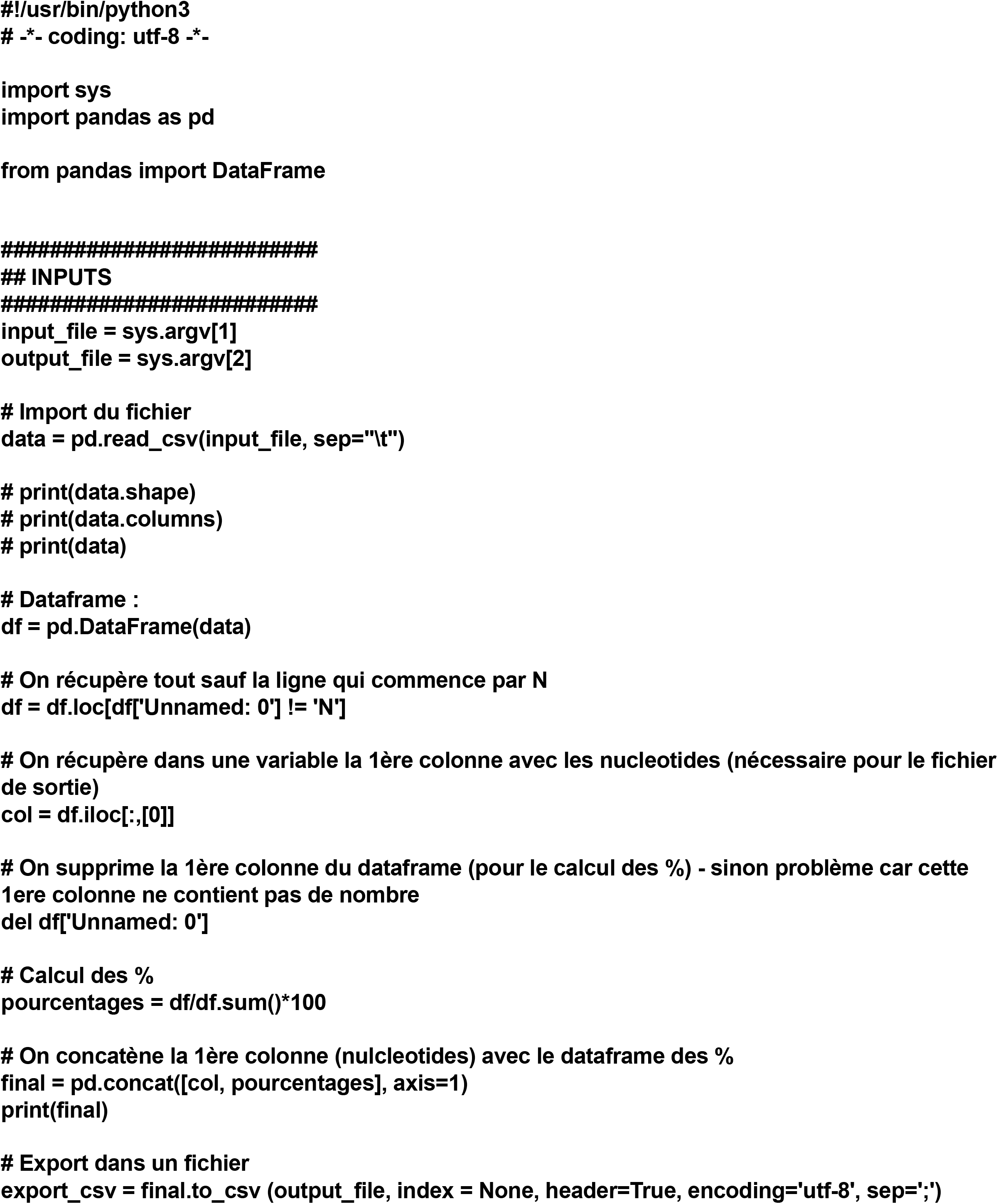
Script (Python) for off-targets analysis.

**Supplementary Figure 3:**
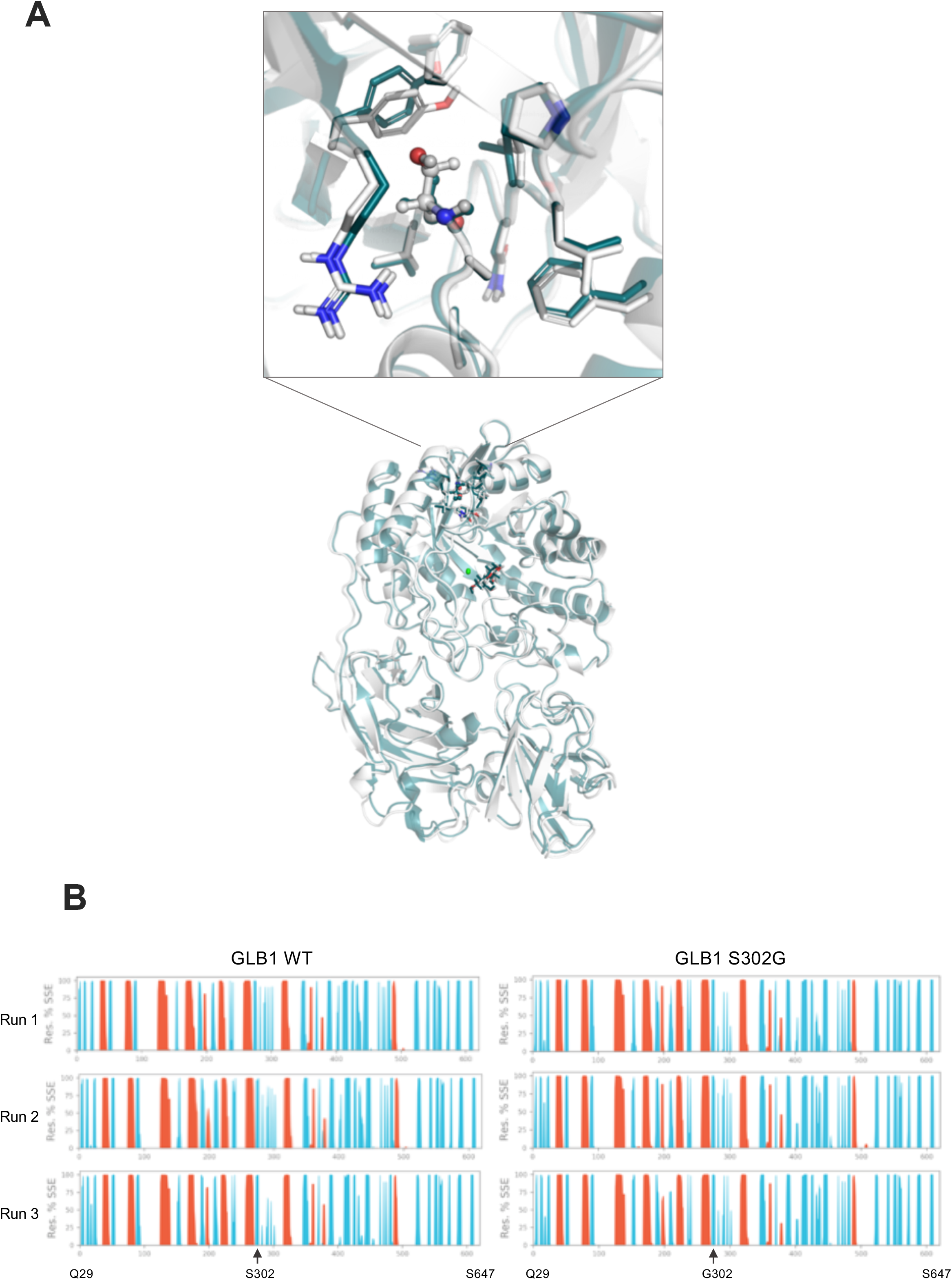
ABE bystander edit and 3D structure predictions. (**A**) Overlay of the energy minimized structures of GLB1 wild type (white) and S302G (teal). (**B**) Analysis of the trajectories (secondary of tertiary structure of GLB1) of three independent molecular dynamics (MD) simulations (3 × 100 ns).

## References

1. Saha, K. et al. The NIH Somatic Cell Genome Editing program. Nature 592, 195–204 (2021).

2. Doudna, J. A. The promise and challenge of therapeutic genome editing. Nature 578, 229–236 (2020).

3. Anzalone, A. V, Koblan, L. W. & Liu, D. R. Genome editing with CRISPR-Cas nucleases, base editors, transposases and prime editors. Nat. Biotechnol. 38, 824–844 (2020).

4. Newby, G. A. & Liu, D. R. In vivo somatic cell base editing and prime editing. Mol. Ther. 29, 3107–3124 (2021).

5. Newby, G. A. et al. Base editing of haematopoietic stem cells rescues sickle cell disease in mice. Nature 595, 295–302 (2021).

6. Krishnamurthy, S. et al. Functional correction of CFTR mutations in human airway epithelial cells using adenine base editors. Nucleic Acids Res. 49, 10558–10572 (2021).

7. Banskota, S. et al. Engineered virus-like particles for efficient in vivo delivery of therapeutic proteins. Cell 185, 250-265.e16 (2022).

8. Koblan, L. W. et al. In vivo base editing rescues Hutchinson-Gilford progeria syndrome in mice. Nature 589, 608–614 (2021).

9. Musunuru, K. et al. In vivo CRISPR base editing of PCSK9 durably lowers cholesterol in primates. Nature 593, 429–434 (2021).

10. Kleinstiver, B. P. et al. High-fidelity CRISPR-Cas9 nucleases with no detectable genome-wide off-target effects. Nature 529, 490–5 (2016).

11. Boutin, J. et al. ON-Target Adverse Events of CRISPR-Cas9 Nuclease: More Chaotic than Expected. Cris. J. 5, 19–30 (2022).

12. Gaudelli, N. M. et al. Directed evolution of adenine base editors with increased activity and therapeutic application. Nat. Biotechnol. 38, 892–900 (2020).

13. Rees, H. A., Minella, A. C., Burnett, C. A., Komor, A. C. & Gaudelli, N. M. CRISPR-derived genome editing therapies: Progress from bench to bedside. Mol. Ther. 29, 3125–3139 (2021).

14. Komor, A. C., Kim, Y. B., Packer, M. S., Zuris, J. A. & Liu, D. R. Programmable editing of a target base in genomic DNA without double-stranded DNA cleavage. Nature 533, 420– 4 (2016).

15. Gaudelli, N. M. et al. Programmable base editing of A•T to G•C in genomic DNA without DNA cleavage. Nature 551, 464–471 (2017).

16. Rees, H. A. et al. Improving the DNA specificity and applicability of base editing through protein engineering and protein delivery. Nat. Commun. 8, 15790 (2017).

17. Walton, R. T., Christie, K. A., Whittaker, M. N. & Kleinstiver, B. P. Unconstrained genome targeting with near-PAMless engineered CRISPR-Cas9 variants. Science 368, 290–296 (2020).

18. Huang, T. P., Newby, G. A. & Liu, D. R. Precision genome editing using cytosine and adenine base editors in mammalian cells. Nat. Protoc. 16, 1089–1128 (2021).

19. Nicoli, E.-R. et al. GM1 Gangliosidosis-A Mini-Review. Front. Genet. 12, 734878 (2021).

20. Lang, F. M., Korner, P., Harnett, M., Karunakara, A. & Tifft, C. J. The natural history of Type 1 infantile GM1 gangliosidosis: A literature-based meta-analysis. Mol. Genet. Metab. 129, 228–235 (2020).

21. McCurdy, V. J. et al. Sustained normalization of neurological disease after intracranial gene therapy in a feline model. Sci. Transl. Med. 6, 231ra48 (2014).

22. Hinderer, C., Nosratbakhsh, B., Katz, N. & Wilson, J. M. A Single Injection of an Optimized Adeno-Associated Viral Vector into Cerebrospinal Fluid Corrects Neurological Disease in a Murine Model of GM1 Gangliosidosis. Hum. Gene Ther. 31, 1169–1177 (2020).

23. Levy, J. M. et al. Cytosine and adenine base editing of the brain, liver, retina, heart and skeletal muscle of mice via adeno-associated viruses. Nat. Biomed. Eng. 4, 97–110 (2020).

24. Li, H. & Durbin, R. Fast and accurate short read alignment with Burrows-Wheeler transform. Bioinformatics 25, 1754–60 (2009).

25. Van der Auwera, G. A. et al. From FastQ data to high confidence variant calls: the Genome Analysis Toolkit best practices pipeline. Curr. Protoc. Bioinforma. 43, 11.10.1-11.10.33 (2013).

26. Wang, K., Li, M. & Hakonarson, H. ANNOVAR: functional annotation of genetic variants from high-throughput sequencing data. Nucleic Acids Res. 38, e164 (2010).

27. Liu, X., Wu, C., Li, C. & Boerwinkle, E. dbNSFP v3.0: A One-Stop Database of Functional Predictions and Annotations for Human Nonsynonymous and Splice-Site SNVs. Hum. Mutat. 37, 235–41 (2016).

28. Ho, M. W. & O’Brien, J. S. Hurler’s syndrome: deficiency of a specific beta galactosidase isoenzyme. Science 165, 611–3 (1969).

29. Kleinstiver, B. P. et al. Engineered CRISPR-Cas9 nucleases with altered PAM specificities. Nature 523, 481–5 (2015).

30. Haeussler, M. et al. Evaluation of off-target and on-target scoring algorithms and integration into the guide RNA selection tool CRISPOR. Genome Biol. 17, 148 (2016).

31. Doench, J. G. et al. Optimized sgRNA design to maximize activity and minimize off-target effects of CRISPR-Cas9. Nat. Biotechnol. 34, 184–191 (2016).

32. Clement, K. et al. CRISPResso2 provides accurate and rapid genome editing sequence analysis. Nat. Biotechnol. 37, 224–226 (2019).

33. Morrone, A. et al. Insertion of a T next to the donor splice site of intron 1 causes aberrantly spliced mRNA in a case of infantile GM1-gangliosidosis. Hum. Mutat. 3, 112– 20 (1994).

34. Anzalone, A. V. et al. Search-and-replace genome editing without double-strand breaks or donor DNA. Nature (2019) doi:10.1038/s41586-019-1711-4.

35. Finkel, R. S. et al. Treatment of infantile-onset spinal muscular atrophy with nusinersen: a phase 2, open-label, dose-escalation study. Lancet 388, 3017–3026 (2016).

36. Akcakaya, P. et al. In vivo CRISPR editing with no detectable genome-wide off-target mutations. Nature 561, 416–419 (2018).

37. Yeh, W.-H., Chiang, H., Rees, H. A., Edge, A. S. B. & Liu, D. R. In vivo base editing of post-mitotic sensory cells. Nat. Commun. 9, 2184 (2018).

38. Chandler, R. J., Sands, M. S. & Venditti, C. P. Recombinant Adeno-Associated Viral Integration and Genotoxicity: Insights from Animal Models. Hum. Gene Ther. 28, 314– 322 (2017).

39. Mangeot, P. E. et al. Genome editing in primary cells and in vivo using viral-derived Nanoblades loaded with Cas9-sgRNA ribonucleoproteins. Nat. Commun. 10, 45 (2019).

